# Resistome-based machine learning source attribution of a superbug

**DOI:** 10.64898/2026.11.16.766933

**Authors:** Carlos-Francisco Méndez-Cruz, David Vargas-Peralta, Luis A. Rendón-Bautista, Santiago Castillo-Ramírez

**Affiliations:** Programa de Genómica Computacional, Centro de Ciencias Genómicas, Universidad Nacional Autónoma de México, Cuernavaca, México; Programa de Genómica Evolutiva, Centro de Ciencias Genómicas, Universidad Nacional Autónoma de México, Cuernavaca, México

**Author notes:** **Corresponding Authors:** Santiago Castillo-Ramírez, Carlos-Francisco Méndez-Cruz.

**Keywords:** Source attribution, Machine Learning, *A. baumannii*, resistome

## Abstract

Many bacterial human pathogens infect not only humans but also other species, and source attribution has been very useful for studying transmission among the different sources. Yet, critical pathogens such as *Acinetobacter baumannii* have not been studied using source attribution approaches. Here, we conducted machine source attribution using the resistome of almost 1200 *A. baumannii* genomes from four sources: humans, companion animals, soil, and white storks. Of note, no other study has used the resistome as input data before. First, we selected the best model among 168 parameter combinations, which was Random Forest, with 5-fold cross-validation and chi2 for feature selection. The model shows a reasonably good accuracy with an F1 weighted score = 0.744, yet class accuracy was dependent on the source. Whereas for companion animals and humans, 93% and 86% of the predictions were correct, respectively; the other two categories (soil and white stork) show correct predictions below 50%. Yet, a core phylogeny and ST assignment reveal that the wrong predictions are associated with multi-host lineages. Finally, we identify the most important ARGs contributing to each class prediction and, in the case of companion animals, *tet* genes, which confer resistance to tetracycline, were among the top contributors. Our study shows the resistome’s potential for developing models to predict source associations. More broadly, this study shows how integrating machine learning and phylogenomics can help us understand multi-host transmission dynamics.

**Importance:** Many bacterial human pathogens infect not only humans, and source attribution has been used for studying transmission among different sources. Yet, critical pathogens such as *Acinetobacter baumannii* have not been studied using source attribution approaches. Here, we conducted machine source attribution using the resistome of almost 1200 *A. baumannii* genomes from humans, companion animals, soil, and white storks. The model shows a reasonably good overall accuracy, yet class accuracy was dependent on the source. Whereas companion animals and humans had high accuracy, the other two categories (soil and white stork) did not. Yet, a phylogeny and ST assignment reveal that the poor predictions are associated with multi-host lineages. Finally, we identify the most important ARGs contributing to each class prediction. This study shows the resistome’s potential for developing models to predict source associations

## Main text

From a public health standpoint, it is very relevant to understand sources and patterns of dissemination of the most important human pathogens. In this regard, source attribution has been very useful in identifying sources and transmission of notable human pathogens, such as *Campylobacter jejuni, Escherichia coli*, or *Salmonella enterica* [1-3]. Yet, there are still critical pathogens, such as *Acinetobacter baumannii*, for which no source attribution strategies have been conducted. Although *A. baumannii* is usually considered a human pathogen, research over the last decade has shown that *A. baumannii* is found in various sources, including animals, plants, and aquatic environments [4, 5]. A key clinical feature of this species is its highly dynamic resistome [6], with many antibiotic resistance genes (ARGs). In a global multi-source study (covering more than 90 different types), we recently showed that there is frequent transmission between humans and many different non-human sources [7]. Here, we revised that dataset to select a set of genomes from different sources to conduct the first study on source attribution for this species using machine learning (ML). We aimed to show that resistome-based source attribution is possible in *A. baumannii*. To have reliable results, we focused only on the most frequent sources. From the global multi-source study [7], we kept 1190 genomes for downstream analyses (see Supplementary Table 1). These genomes include 4 different sources (see Figure 1A): companion animals (cats and dogs), humans, soil, and white storks. The resistome of the isolates was inferred as we did for the original study [7]; see Supplementary File 1 for all technical details. We chose to use the ARGs (the resistome) as input data for our source attribution strategy for several reasons. First, *A. baumannii* from different sources seems to have different ARGs [8-10]. Second, at least for grass, cattle, and pigs, the profile of ARGs matched well with the phylogeny [8, 9]. Finally, given that we know a lot about the ARGs, the interpretability of the source attribution results can be more feasible.

**Figure 1.**
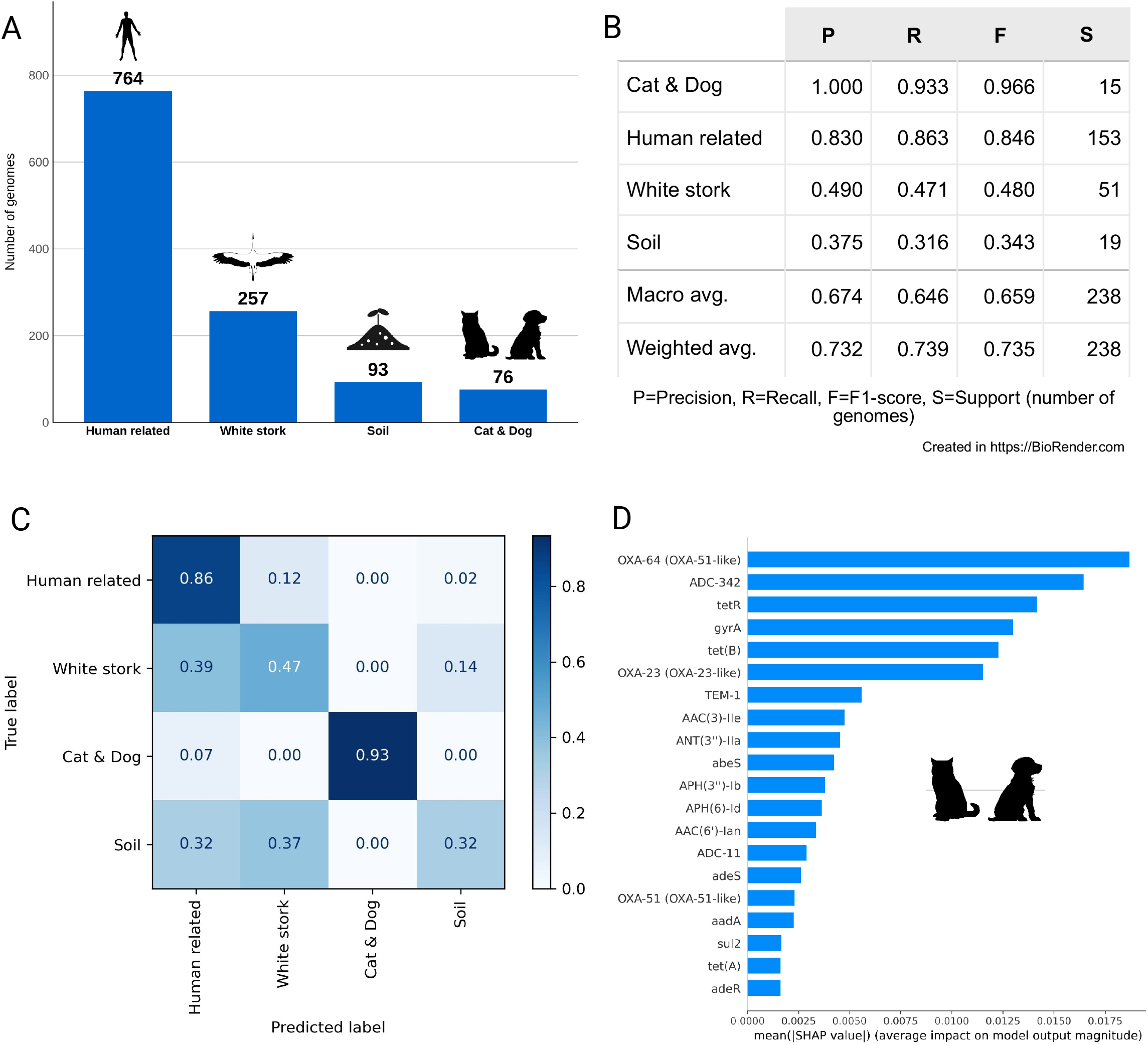
Input data, accuracy of the best model, and model interpretability. Panel A: the different sources considered for the study; the bars give the number of isolates (genomes) per source. Panel B: classification report with the testing data set. Panel C: confusion matrix normalized by row with the testing data set. Panel D: SHAP-based ARG importance ranking for cats & dogs source; the panel shows the top 20 ARGs influencing the prediction.

Considering the machine learning model, we split the data into a training data set (80%) and a testing data set (20%), but we paid attention to stratifying the training-testing splits according to source frequency (see Supplementary Figure 1). To develop the best model, we considered 4 algorithms (KNN, SVM, RF, and XGB), two k-fold cross-validation schemes, and different feature selection and oversampling strategies. Details are provided in Supplementary File 1 and Supplementary Figure 2. The best model was RF, with a 5-fold cross-validation, chi2 for feature selection, and with no oversampling correction (see Supplementary Figure 2). This model had an F1-weighted score = 0.744, and notably, with the test data set it had a very minor reduction in the F1-weighted score = 0.735. Thus, at a general level (across all classes), the model showed a reasonably good accuracy. Analysing the classification report for the different sources (see Figure 1B), we found accuracy was dependent on the source. Whereas for 2 categories (cat_dogs and human), we noted very good F1 score values (0.966 and 0.846, respectively), for the other two (soil and white stork), the accuracy was low, with F1 scores below 0.5. It is important to mention that the number of genomes per category might not explain the class accuracy variation, as the category with the fewest genomes (cat+dogs) and the category with the most genomes (human) have the highest F1 scores. To properly understand the model’s poor predictions (Figure 1C), we analyse the confusion matrix in the context of a core genome phylogeny and also consider ST assignment. Figure 2 shows a core phylogeny for all the isolates and highlights the misclassification cases (external black dots and colored bars in the internal ring). We observed that many misclassification cases are located in clades having isolates from several sources. We now provide some concrete examples. The blue arrows show some cases of human isolates misclassified as white storks, yet these cases fall in clades with both human and white stork isolates. The purple circle shows a cluster of several misclassifications in a clade. For instance, there are 2 white stork isolates misclassified as human, one human isolate misclassified as white stork, and even a soil isolate misclassified as white stork. However, that clade actually contains all the sources except for companion animals. It is important to note that in all these apparent misclassification cases, the isolate “wrongly classified” is very closely related to isolates with the same source as the wrong classification. The ST assignment further reinforces this. Given that Pasteur MLST is more accurate than Oxford MLST [11], all the STs mentioned below are as per the former. For instance, 4 white stork isolates misclassified (2 as human and 2 as soil) were assigned to STs (ST155, ST199, ST1079) already described in humans. Notably, ST155 has been described in humans but also in bovine cattle [12]. Also, in the only wrong prediction for the cat+dogs category, we noted that this isolate belongs to ST1, an ST previously associated with cat, dog, and human isolates. Thus, the apparent misclassification cases point to isolates belonging to multi-host lineages, a real biological phenomenon. It is relevant to mention that in the original multi-host source study [7], multi-host lineages were a common occurrence. Finally, to identify the most relevant ARGs across the classes (sources), we used the SHapley Additive exPlanations (SHAP) analysis (see Figure 1D and Supplementary Figure 3). In the case of cats and dogs, we mention some relevant ARGs. For instance, three *tet* genes [tet(A), tet(B), and tetR] were found, with two of them being ranked second and fifth in terms of their SHAP values. Of note, these genes are related to tetracycline resistance; here, it is worth mentioning that doxycycline (a member of the tetracycline family) is extensively prescribed in veterinary medicine to treat bacterial infections in pets due to its broad-spectrum antibiotic activity [13]. Another example is OXA-64, a variant of the intrinsic Oxacillinase 51-like family, ranked first. Worth paying attention is that OXA-64 is associated with ST25, which is reported in both cats and dogs [14]. We also found OXA-23 (*blaOXA-23*), which is the most commonly acquired OXA. Of note, this ARG has been previously noted in cats and dogs [7, 15]. Supplementary Figure 3 provides the SHAP analysis details for the other sources.

**Figure 2.**
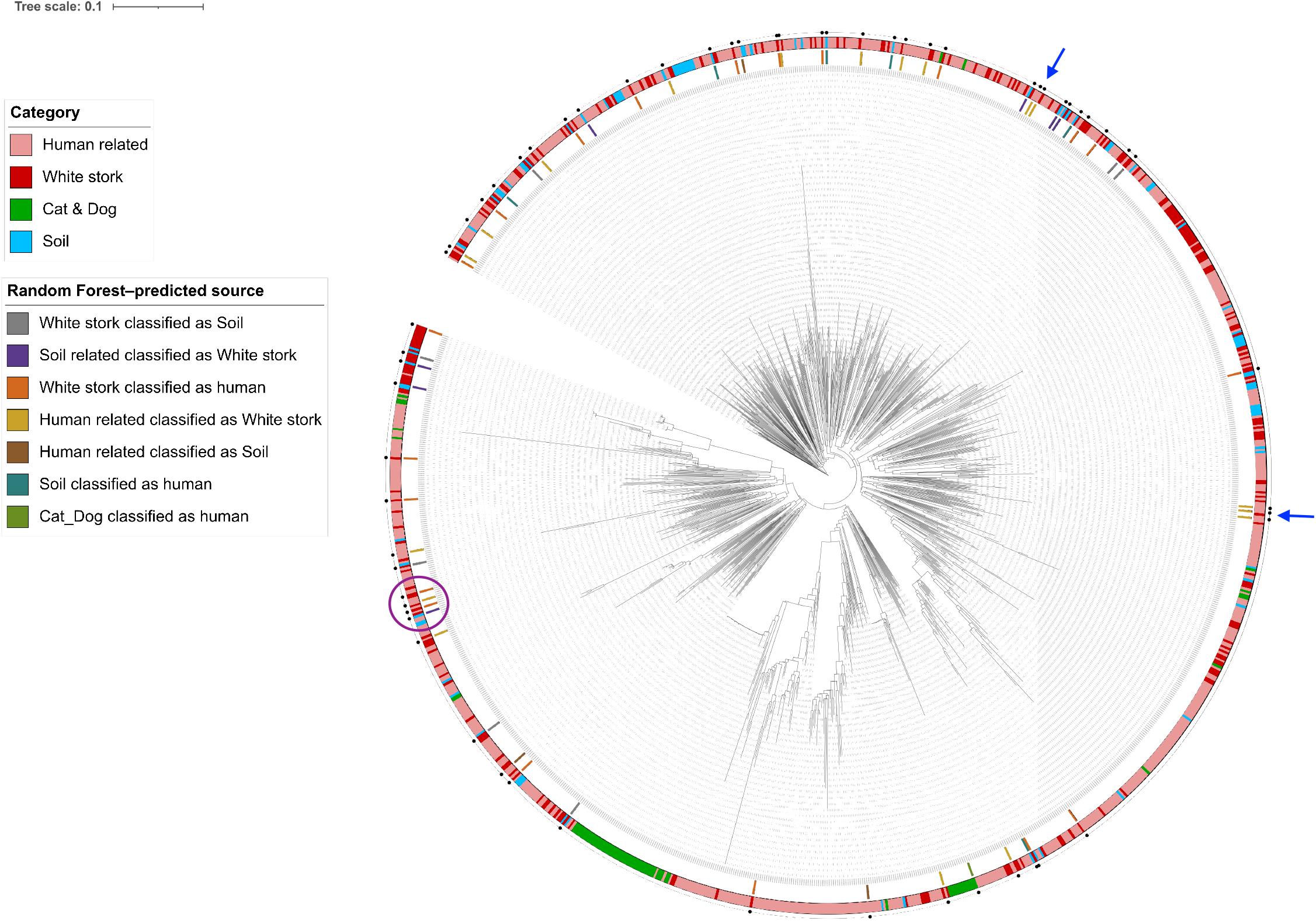
Core genome phylogeny. Core-genome phylogeny of the genomes considered in this study. The external ring gives the source of the isolates, whereas the internal ring highlights the “wrong” predictions by the model. The blue and purple rings denote some cases of wrong classifications. The scale gives the number of substitutions per site.

In conclusion, this study shows that source attribution can be implemented for *A. baumannii*. We think that this work is a proof of principle that the resistome can be used to carry out source attribution in many bacteria with a decent number of ARGs. Yet, we acknowledged that using more genes, without biasing them to any particular function, such as the whole accessory genome, would likely have more resolution and might produce more accurate results for non-human categories. In that regard, we are already running preliminary analyses considering the whole pangenome of the species. Nonetheless, it is clear that this study is bound to be a reference point for future studies conducting source attribution for *A. baumannii*. Finally, our study also shows how integrating phylogenies and ST assignments can help make sense of the ML source attribution predictions.

## Data Sharing

Supplementary Table 1 lists all the genomes, with their accession numbers, used for this study.

## Funding Declaration

David Vargas-Peralta is a doctoral student in the Posgrado en Ciencias Biológicas, and Luis A. Rendón Bautista is a doctoral student in the Doctorado en Ciencias Bioquímicas, both at Universidad Nacional Autónoma de México (UNAM). They received scholarships from Secretaría de Ciencias, Humanidades, Tecnología e Innovación (SECIHTI) under scholarship numbers 816227 and 928282, respectively. The students gratefully acknowledge the financial support provided.

## Authors contributions

SCR conceived the project. CFMC and SCR coordinated the project. CFMC and DVP curated the data collection. CFMC conducted most of the machine learning analyses. DVP ran the phylogeny and created the cartoons in Figure 1. LARB conducted some of the machine learning analyses. CFMC and LARB wrote the supplementary methods. SCR drafted the article, and all the authors approved the final version of the article.

## Conflicts of interest

The authors declare that there are no conflicts of interest.

## SUPPLEMENTARY FIGURES

**Supplementary File 1**

Description of the methodology used for this study.

**Supplementary Figure 1**

Stratified random split the data into a training data set (80%) and a testing data set (20%).

**Supplementary Figure 2**

Finding the best model

To develop the best model, we considered 4 algorithms (KNN, SVM, RF, and XGB), two k-fold cross-validation schemes, and different feature selection and oversampling strategies. The best model (RF) was evaluated with the testing data set.

**Supplementary Figure 3**

SHAP values for the top ARGs for three categories (human, soil, and white storks).

## SUPPLEMENTARY TABLES

**Supplementary Table 1**

List of all the genomes considered in this study along with their metadata.

**Supplementary Table 2**

Resistome prediction for all the genomes considered.

